# Macrophages Promote Aortic Valve Cell Calcification Through STAT3 Splicing

**DOI:** 10.1101/2020.01.24.919001

**Authors:** Michael A. Raddatz, Tessa M. Huffstater, Matthew R. Bersi, Bradley I. Reinfeld, Matthew Z. Madden, Sabrina E. Booton, W. Kimryn Rathmell, Jeffrey C. Rathmell, Brian R. Lindman, Meena S. Madhur, W. David Merryman

## Abstract

**Objective:** Macrophages have been described in calcific aortic valve disease, but it is unclear if they promote or counteract calcification. We aimed to determine how macrophages are involved in calcification using the *Notch1*^+/-^ model of calcific aortic valve disease.

**Approach and Results:** Macrophages in wild-type and *Notch1*^+/-^ murine aortic valves were characterized by flow cytometry. Macrophages in *Notch1*^+/-^ aortic valves had increased expression of MHCII. We then used bone marrow transplants to test if differences in *Notch1*^+/-^ macrophages drive disease. *Notch1*^+/-^ mice had increased valve thickness, macrophage infiltration, and M1-like macrophage polarization regardless of transplanted bone marrow genotype. *In vitro* approaches confirm that *Notch1*^+/-^ aortic valve cells promote macrophage invasion as quantified by migration index and M1-like polarization quantified by Ly6C and CCR2 positivity regardless of macrophage genotype. Finally, we found that macrophage interaction with aortic valve cells promotes osteogenic, but not dystrophic, calcification by decreasing abundance of the STAT3β isoform.

**Conclusions:** This study reveals that *Notch1*^+/-^ aortic valve disease involves increased macrophage recruitment and polarization driven by altered aortic valve cell secretion, and that increased macrophage recruitment promotes osteogenic calcification through STAT3 splicing changes. Further investigation of STAT3 and macrophage-driven inflammation as therapeutic targets in calcific aortic valve disease is warranted.

**Graphical Abstract:** 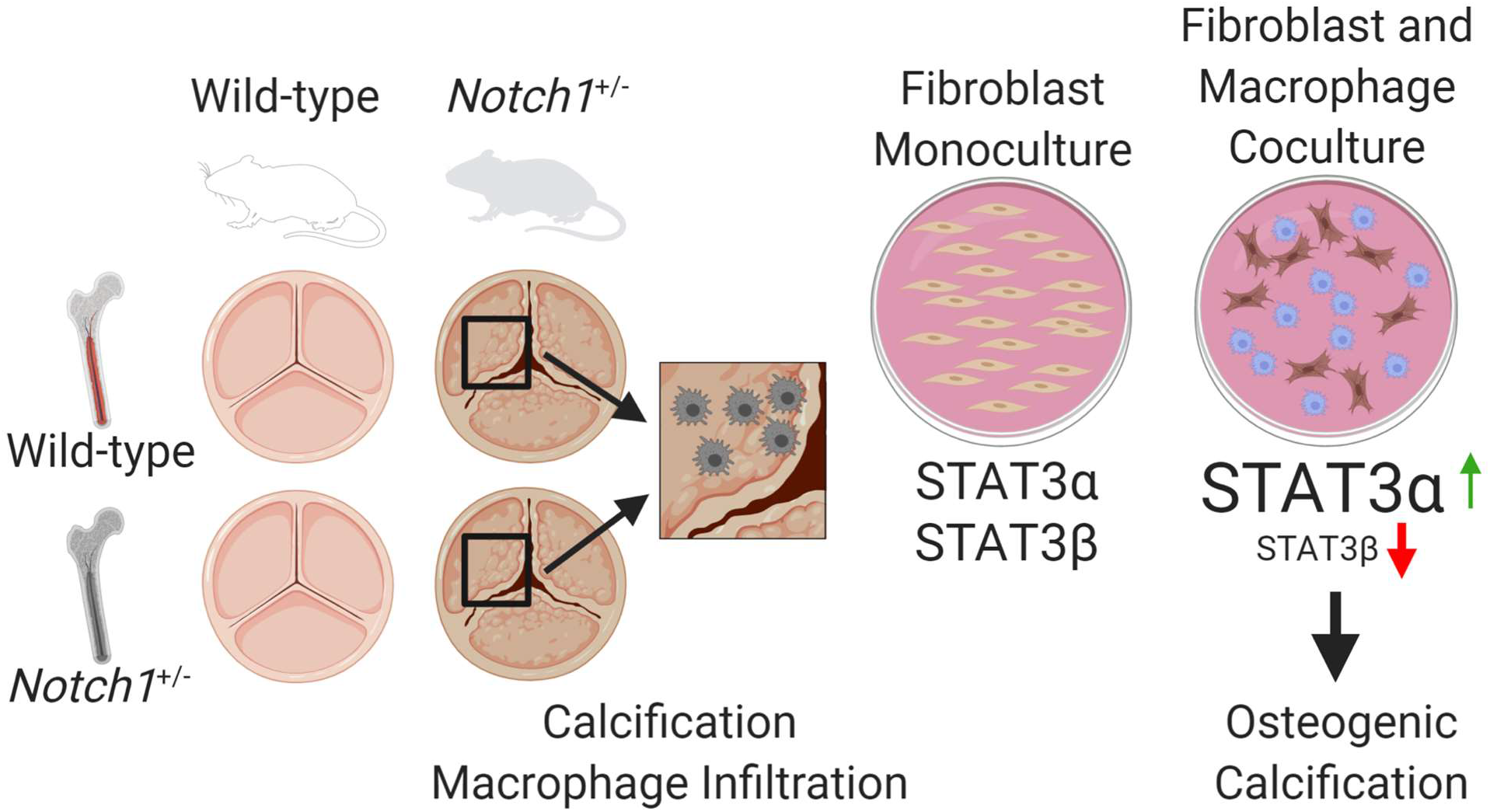

## Introduction

Calcific aortic valve disease (CAVD) affects one in four people over 65 years of age and is the primary cause of aortic stenosis.^1, 2^ This prevalent and insidious disease inevitably leads to surgical or transcatheter replacement of the valve, as there are currently no pharmaceutical treatments. Understanding cellular and molecular pathophysiology may lead to pharmaceutical approaches for patients who are not optimal surgical candidates or prevent prosthetic valve recalcification.

For the majority of the field’s existence, CAVD studies have focused on aortic valve interstitial cells (AVICs),^3^ yet no successful pharmacological strategies have emerged from this approach.^1^ Recent studies have shown that inflammation and immunomodulation may play a key role in determining the calcification potential of these cells,^4–6^ suggesting that immune signaling may be a viable target for therapeutic intervention. Immune cells, and specifically macrophages, are linked to CAVD,^7^ and up to 10-15% of cells in the healthy valve are CD45^+^, a hematopoietic lineage marker.^8, 9^ These cells are primarily major histocompatibility complex II^+^ (MHCII^+^) macrophages.^8, 10^ M1-like macrophages, of which MHCII positivity is a marker, are more metabolically active, direct a proinflammatory immune response, and are further increased in CAVD.^5, 11, 12^ As part of this proinflammatory response, these macrophages secrete interleukin-6 (IL-6) and tumor necrosis factor α (TNFα), both of which promote calcification of AVICs.^5^ However, macrophage depletion with liposomal clodronate increases disease as measured by aortic valve (AV) thickening in mice.^13^ It is unclear if macrophages drive CAVD, inhibit CAVD, or respond to calcification.

Moving from cellular to molecular inflammation, STAT3 signaling is linked to both the activity of the osteogenic transcription factor RUNX2 and fibrotic inflammation in the heart.^14–18^ These activities reflect the two primary pathways of AV calcification: osteogenic and dystrophic calcification, respectively.^19, 20^ Additionally, single nucleotide polymorphisms in IL-6 receptor (a major contributor to STAT3 activation) are associated with decreased severity of CAVD,^21^ and transforming growth factor beta 1 (TGF-β1, another direct activator of STAT3) is increased in CAVD and leads to calcification of AV cells *in vitro*.^22, 23^ Adding to the evidence, Tsai, et al. reported increased STAT3 phosphorylation in human CAVD.^24^ This confluence of findings suggests that STAT3-mediated inflammation, potentially driven by macrophage-secreted factors, may promote CAVD and serve as a pharmacological target.

In order to determine the role of immune cells in originating CAVD, we utilized the *Notch1*^+/-^ murine CAVD model; human families with *NOTCH1* mutations have increased incidence of both CAVD and congenital bicuspid AV disease.^25^ Mice with *Notch1* haploinsufficiency have increased AV calcification,^26, 27^ while AVICs with a *Notch1* mutation have increased calcification potential *in vitro*.^28^ Interestingly, *NOTCH1* has long been known to play a significant role in the differentiation and maturation of hematopoietic lineages—including specific inhibition of myeloid development—thus highlighting the potential for haploinsufficiency to affect valve calcification through infiltrating macrophages.^29–31^ After assaying macrophage phenotypes in the *Notch1*^+/-^ model, we utilized bone marrow transplants and *in vitro* coculture models to assess the contribution of *Notch1*^+/-^ immune cells to calcification. Finally, we assessed and manipulated STAT3 activity using overexpression plasmids and phosphorylation blockade to investigate the contribution of macrophage-mediated changes in STAT3 to AVIC calcification. We found that *Notch1*^+/-^ AVICs increase recruitment and inflammatory maturation of macrophages, and that macrophages promote AVIC calcification in part through STAT3 signaling.

## Methods

### Animal Studies

All animal experiments used C57BL/6J *mus musculus* animals. Bone marrow transplant experiments used both sexes, while experiments with less than 8 mice per group used male mice only. In total, 114 mice were used for this study. All experimental protocols were approved by the Institutional Animal Care and Use Committee at Vanderbilt University.

### Flow cytometry

AVs were isolated from littermate wild-type and *Notch1*^+/-^ mice. Cells were isolated from the AV by nine, seven-minute collagenase digestions at 37°C. After each digestion, the supernatant was removed and diluted into FC buffer (PBS, 3% FBS). The cell pellet was then subjected to red blood cell lysis buffer for 5 minutes before quenching with FC buffer. For *in vitro* assays, BMMs and/or AVICs were lifted with Accutase. Cells were then blocked in Fc Block for 10 minutes at room temperature before staining with conjugated antibody for 30 minutes at 4°C (Please see the Major Resources Table in the Supplemental Materials).

### Bone marrow transplants

8- to 12-week-old wild-type or *Notch1*^+/-^ C57BL/6J mice were given a split 12 Gy dose of radiation from a Cs^137^ source followed by retro-orbital administration of 3×10^6^ bone marrow cells isolated from a sex-matched wild-type or *Notch1*^+/-^ donor. Mice were allowed 6 weeks for bone marrow reconstitution and aged on high-fat diet. After 6 months, mice were euthanized and bone marrow and AVs were isolated. Bone marrow was digested in rat tail lysis buffer overnight and genotyped for *Notch1* and the *Notch1*^del^ cassette using polymerase chain reaction (PCR) to confirm successful transplants.

### Histology and Immunofluorescence

Murine AVs were frozen in OCT and sectioned at 10 µm thickness. Von Kossa staining was performed by incubating with 3% Ag_2_NO_3_ for 40 minutes under a UV lamp, followed by incubation with 5% sodium thiosulfate for 5 minutes. Slides were counterstained with Nuclear Fast Red. Leaflet thickness was measured using a semi-automated MATLAB script to calculate leaflet area and divide by leaflet length to give average leaflet width. Slides were fixed and permeabilized with 10% formalin and 0.1% Triton-X for 15 minutes, followed by epitope blockade for 1 hour with 1% BSA in PBS. Primary antibody staining was performed in blocking solution overnight at 4°C. When applicable, secondary antibody staining was performed for 1 hour at room temperature. Slides were mounted in ProLong Gold with DAPI and imaged at 4X magnification. CD68^+^ and MHCII^+^ macrophages were counted manually and normalized to the area of a DAPI mask of the leaflet of interest, giving macrophages/mm^2^.

### Aortic valve interstitial cells

AVICs were isolated from wild-type or *Notch1*^+/-^ C57BL/6J mice as previously described.^32^ Briefly, AVs were digested in 2 mg/mL collagenase in HBSS for 30 minutes at room temperature and then placed in DMEM supplemented with 10% FBS, 1% penicillin/streptomycin (pen/strep), and 10 μg/mL recombinant murine interferon-γ to induce activation of the simian virus 40 T antigen. Cells were allowed to adhere to 0.1% gelatin-coated six-well tissue culture-treated plates and expanded. To allow for sustained immortal growth, cells were cultured at 33°C and 5% CO_2_ when not plated for experiments. At least 12 hours prior to experiments, AVICs were incubated at 37°C and 5% CO_2_ in DMEM supplemented with 10% FBS and 1% pen/strep (complete media), wherein the immortalization element degrades due to temperature changes. AVICs were seeded at 20,000 cells/cm^2^ in all experiments unless otherwise noted.

### Bone marrow-derived macrophages

Macrophages (BMMs) were generated from the bone marrow of wild-type or *Notch1*^+/-^ C57BL/6J mice using M-CSF.^33^ BMM generation was verified by flow cytometry for CD11b and F4/80 (Supp Fig I). All experiments post-differentiation were carried out without M-CSF supplementation.

### Coculture design

BMMs and AVICs were seeded at a 1:7 physiologic ratio^8^ in RPMI supplemented with 10% FBS and 1% pen/strep and cultured for 48 hours before harvesting for various experiments. Transwell cocultures were seeded at the same ratio with AVICs seeded on the tissue culture-treated plate and BMMs seeded on a 0.4 μm-pore Transwell insert (Corning, Corning, NY). AVIC monoculture controls for all coculture experiments were also performed in supplemented RPMI.

### Cultured media

Media was harvested from cultures after 24 hours and filtered using 0.45 μm sterile filters before use. In all cultured media experiments, RPMI supplemented with 10% FBS and 1% pen/strep was used.

### Migration assay

Using a modified Transwell migration protocol,^34^ 10,000 BMMs were seeded in 100 μL of uncultured media on an 8 μm-pore Transwell insert and incubated for 10 minutes at 37°C. 600 μL of cultured media was then added to the well below each insert and cells were allowed to migrate for 3 hours. Transwell inserts were then removed and fixed in 70% ethanol for 10 minutes prior to mounting in ProLong Gold with DAPI. All cells that migrated through the membrane were counted based on visualization of DAPI staining. Migration index was defined as the number of migrated cells divided by the number of migrated cells into a control condition of uncultured media.

### Microarray

The Proteome Profiler Mouse Cytokine Array Kit, Panel A (R&D Systems, Minneapolis, MN) was used per the manufacturer’s instructions. Briefly, protein from either cell lysates or conditioned media was incubated with an antibody mixture and allowed to bind to the patterned membrane overnight. Antibodies were then conjugated and the membrane was imaged using an Odyssey Classic imager (Li-Cor, Lincoln, NE). For each membrane experiment, biological replicates were pooled and one membrane was used per condition. All data reported is the mean of two technical replicate spots.

### Calcific nodule assay

Cultures were treated with 5 ng/mL TGF-β1 for 24 hours followed by 24 hours of cyclic biaxial 10% mechanical strain at 1 Hz on BioFlex plates coated with Pronectin, using a FlexCell 3000 machine (Flexcell, Burlington, NC). Cultures were stained for calcification using Alizarin Red S and calcific nodules were counted in each well.

### Magnetic-activated cell sorting

Cells were lifted with Accutase, incubated for 15 minutes with anti-CD45 MicroBeads (130-110-618, Miltenyi Biotec, Bergisch Gladbach, Germany) to allow for magnetic labeling, and resuspended in MACS buffer (PBS, 0.5% BSA, 2 mM EDTA), followed by positive selection of CD45^+^ BMMs. Futher downstream anaylsis was conducted on the unperturbed CD45^-^ AVICs.

### Quantitative real time-polymerase chain reaction (RT-qPCR)

AVIC mRNA was isolated using Trizol (Life Technologies, Carlsbad, CA) and cDNA libraries were produced using the Superscript IV Reverse Transcriptase kit with oligo(dT) primer (ThermoFisher Scientific, Waltham, MA) as per manufacturer protocols. Quantitative real time polymerase chain reaction for all targets was performed on the CFX-96 Real Time System using iQ SYBR Green Supermix (Bio-Rad, Hercules, CA). Products were confirmed by gel electrophoresis. Gapdh was used as a housekeeping gene. All statistics were performed on untransformed ΔCt values (“gene of interest” Ct – *Gapdh* Ct), but for clarity, gene expression was normalized and displayed as 2^ΔΔCt^.

### Cell proximity analysis

Cocultures were performed on glass coverslips and stained by immunofluorescence for CD68 and either RUNX2 or αSMA. Immunofluorescence staining was performed as described above (see *Histology and Immunofluorescence*). Immunofluorescence images were analyzed using a custom algorithm designed to determine whether the proximity of activated AVICs—as identified by RUNX2 or αSMA staining—to CD68^+^ macrophages is closer or further than expected based on Monte Carlo simulations of random macrophage placement. Additional details are included in the Supplemental Materials (Supp Fig II).

### Western blot

AVICs and human AVs were lysed in RIPA buffer or PBS respectively, supplemented with benzonase, sodium orthovanadate, and protease inhibitor. Lysates were denatured using SB at 100°C for 5 minutes, then 10-15 μg was loaded into 15 cm 10% acrylamide gels and run at 150V for 1 hour and 45 minutes. Membrane transfer was performed at 80V for 1 hour and 45 minutes. Membranes were blocked in TBST + 5% BSA and stained in primary antibody overnight at 4°C. Membranes were then stained with fluorescent secondary antibody and imaged on an Odyssey Classic imager (Li-Cor). Quantification was performed in Image Studio Lite (Li-Cor).

### Human aortic valve samples

AV samples were collected at the time of replacement and separated into involved and uninvolved tissue based on the sample location relative to apparent calcification before being flash frozen in liquid nitrogen and stored at −80°C. Samples were mechanically digested with a bead homogenizer (BioSpec Products, Bartlesville, OK) in PBS supplemented with benzonase, sodium orthovanadate, and protease inhibitor. Written informed consent was obtained from patients and tissue sample collection was approved by the institutional review board at Washington University in St. Louis.

### Plasmid transfection

Prior to transfection, AVICs were serum-starved in 1 mL of DMEM with 1% FBS overnight in 12-well plates. Lipofectamine 2000 (ThermoFisher) and concentrated STAT3α, STAT3β, or vector control plasmids (Genscript, Piscataway, NJ) were diluted in Opti-MEM media (ThermoFisher) and allowed to create DNA-lipid complexes for 20 minutes. Next, 200 μL of Opti-MEM containing 4 μL of Lipofectamine and 1 μg of plasmid DNA was added to each well. After 4 hours, media was replaced with complete media. In coculture models, macrophages were added 24 hours after transfection initiation, and in all experiments AVICs were harvested at 48 hours.

### Micropipette aspiration

Micropipette aspiration was used to determine the elastic modulus of AVICs as reported previously.^23, 35–37^ Additional details are included in the Supplemental Materials (Supp Fig III).

### STAT3 blockade

Stattic (MilliporeSigma), a STAT3 tyrosine phosphorylation inhibitor (Y705) was used to block STAT3 activity. Stattic was solubilized in DMSO and added to cells in complete media.

### Statistics

All datapoints are shown throughout the manuscript in addition to mean ± standard error of the mean (s.e.m.) or boxplots signifying median and first and third quartiles for non-normal data. Comparisons between normal data were performed by ANOVA followed by Student’s t-test with Holm-Sidak adjustment for multiple comparisons; non-normal data were analyzed using Kruskal-Wallis or Mann-Whitney *U* test. Murine data were analyzed by aligned rank transformed ANOVA^38^ to allow for two- and three-way non-parametric comparisons. All statistical analyses were performed using the statistical programming language R, version 3.5.2.^39^ The authors declare that all supporting data are available within the article [and its online supplementary files].

## Results

### *Notch1*^+/-^ Aortic Valves Have an Altered Myeloid Profile

We first assessed macrophage phenotypes in the AVs of both wild-type and *Notch1*^+/-^ mice in young adulthood (10-12 weeks), prior to disease onset. In both wild-type and *Notch1*^+/-^ mice, hematopoietic cells make up 5-10% of the AV (Fig 1A). Among hematopoietic cells, there is a majority myeloid fraction (CD11b^+^) that is greater in the *Notch1*^+/-^ valve (Fig 1B) and primarily F4/80^+^ macrophages in both genotypes (Fig 1C). Valvular macrophages are majority CX_3_CR1^high^/Ly6C^-^/CCR2^-^ in both genotypes (Fig 1D-F), but in the *Notch1*^+/-^ valve, macrophages show increased MHCII expression, suggesting an enhanced pro-inflammatory phenotype (Fig 1G). Non-myeloid cell types include CD11b^-^/MHCII^+^ antigen-presenting cells, similar to previous reports (Fig 1H, I).^8^ Simultaneously, BMMs were generated from the same wild-type and *Notch1*^+/-^ mice and no differences were found (Supp Fig IV). In summary, macrophages make up the majority of hematopoietic cells in the AV and are different in the valves of *Notch1*^+/-^ vs wild-type mice.

**Figure 1.**
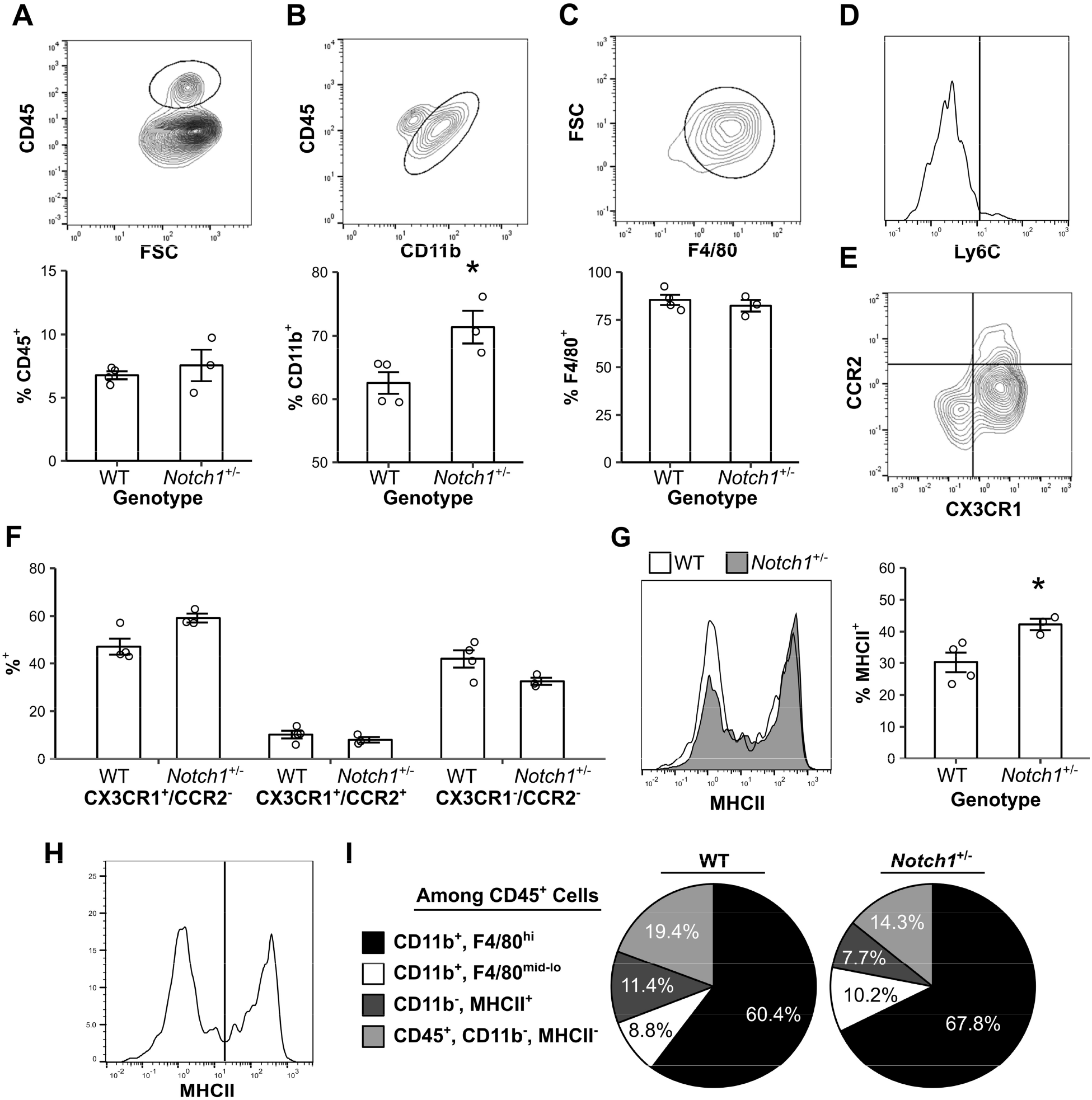
*Notch1*^+/-^ murine valves have increased macrophage polarization. Flow cytometry was performed on both wild-type and *Notch1*^+/-^ aortic valves for CD45 (A), followed by CD11b (B), then F4/80 (C). Macrophages were characterized by Ly6C expression (D), CX3CR1 and CCR2 expression (E, F), and MHCII expression (G, outline = wild-type; gray fill = *Notch1*^+/-^). Non-myeloid hematopoietic cells were grouped by MHCII expression (H), and all cell types plotted by percentage of the CD45^+^ population (I, average of animals from two experiments). Bar plots represent mean ± s.e.m from one of two identical experiments (A-C, F, G). Representative flow plots are of wild-type animals (A-E, H). *P < 0.05 by two-tailed t test. N = biological replicates.

### *Notch1* Haploinsufficiency in AVICs Drives Calcification and Macrophage Recruitment

Considering the increased macrophage infiltration in *Notch1*^+/-^ mice previously reported,^26^ and the altered macrophage phenotypes found above, we performed bone marrow transplants to identify if underlying differences in hematopoietic cells were driving the macrophage changes found in the *Notch1*^+/-^ CAVD model. Wild-type and *Notch1*^+/-^ mice were transplanted with wild-type or *Notch1*^+/-^ bone marrow and aged for 6 months on high-fat diet to allow for disease progression. After aging, body genotype, but not bone marrow genotype, was significantly associated with leaflet thickness (Fig 2A, B) and non-significantly associated with leaflet calcification after von Kossa staining (Fig 2A, C). The same pattern was seen with immunofluorescence staining for CD68^+^ macrophage infiltration (Fig 2D, E). Valves were additionally stained for MHCII to detect differences observed by flow cytometry (Fig 2F, G). Valves of *Notch1*^+/-^ mice have an increased prevalence of MHCII^+^ macrophages and a lesser increase in MHCII^-^ macrophages, leading to a higher MHCII^+^ macrophage fraction (Fig 2H, I, Supp Fig V). Thus, *Notch1* haploinsufficient valve phenotypes are mediated by valvular cells, rather than infiltrating hematopoietic cells.

**Figure 2.**
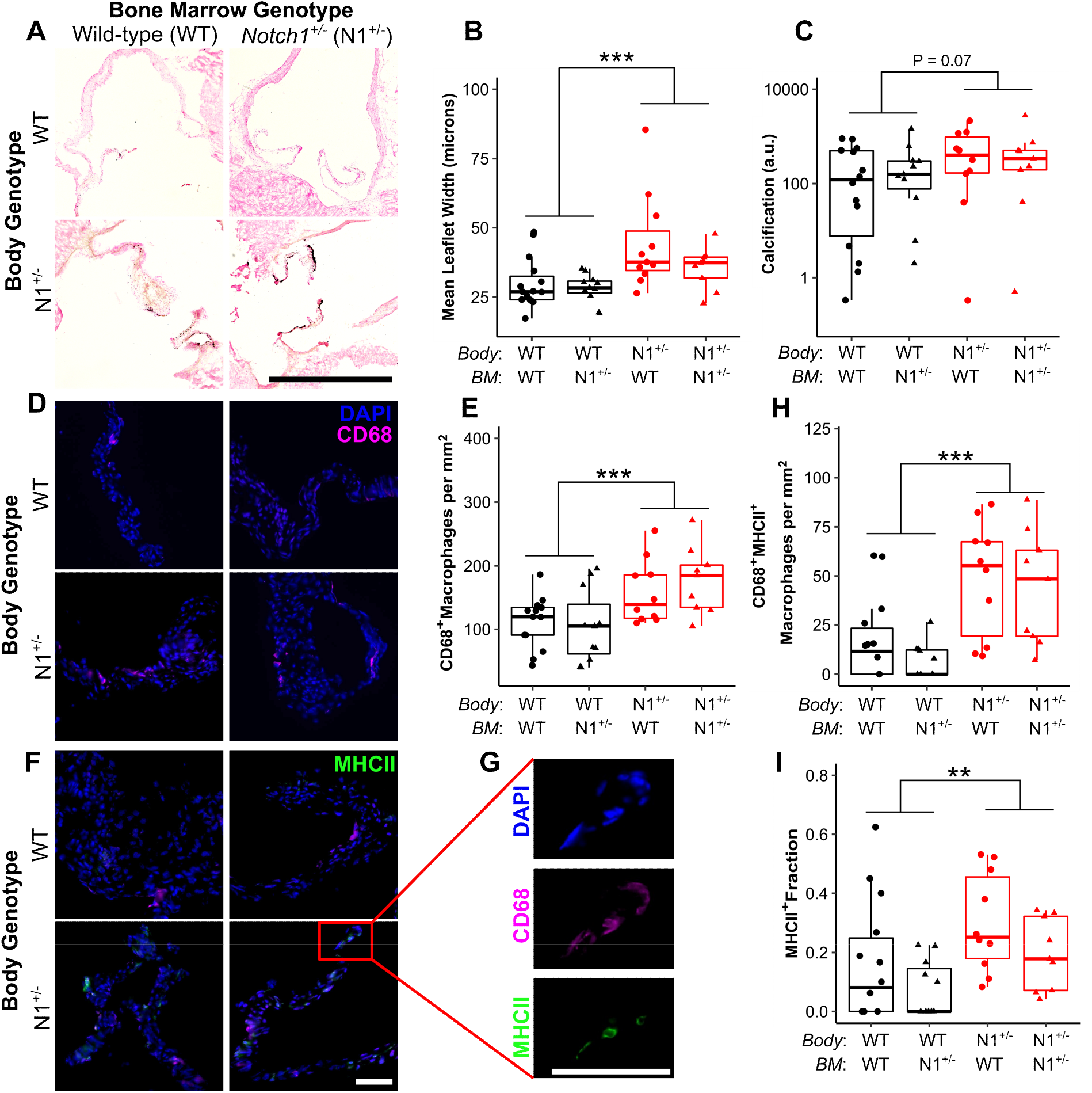
*Notch1*^+/-^ valve cells drive aortic valve calcification and macrophage infiltration and maturation *in vivo*. Following bone marrow transplant, *Notch1*^+/-^ (N1^+/-^) and wild-type (WT) mice were assessed for aortic valve thickness (A, B), calcification (A, C), macrophage infiltration (D, E), and macrophage maturation as measured by MHCII positivity (F, G, H, I). (A) Aortic valves are stained for histology and calcification by von Kossa; scale bar = 1 mm. (B) Leaflet thickness is plotted as mean width across the entire section. (C) Calcification is measured by positive pixels per section. (D, F, G) Aortic valves are stained by immunofluorescence for DAPI (blue), CD68 (pink), and MHCII (green); scale bar = 100 µm. (E, H) Macrophage data is plotted as cells per mm^2^ of tissue. Boxplots represent 25^th^, 50^th^, and 75^th^ percentiles. All data were analyzed by two-way aligned rank transformed ANOVA.^38^ **P < 0.01, ***P < 0.001. N = biological replicates. BM = bone marrow.

### *Notch1*^+/-^ AVICs Drive Calcification and Macrophage Phenotypes *In Vitro*

With *Notch1* haploinsufficiency acting through AVICs but accompanied by a clear difference in macrophage infiltration and phenotype, we used *in vitro* coculture models to explore this relationship. First, we replicated the previous bone marrow transplant experiment *in vitro*. *Notch1*^+/-^ and wild-type AVICs and BMMs were cocultured and assayed for common transcriptional calcification markers. *Notch1* haploinsufficiency altered coculture calcification genes only when carried in the AVICs (Fig 3A-D). Assessing macrophage phenotypes, BMMs cultured with *Notch1*^+/-^ AVICs had increased CCR2 and Ly6C expression, while macrophage genotype had no effect (Fig 3E, F). Additionally, both wild-type and *Notch1*^+/-^ macrophages increase migration towards media cultured by *Notch1*^+/-^ AVICs compared to wild-type AVICs (Fig 3G, Supp Fig VI). Microarray analysis of secreted factors and lysate from *Notch1*^+/-^ and WT AVICs reveal an increase in cytokines that induce M1 differentiation and migration (Fig 3H, Supp Fig VII).^40, 41^

**Figure 3.**
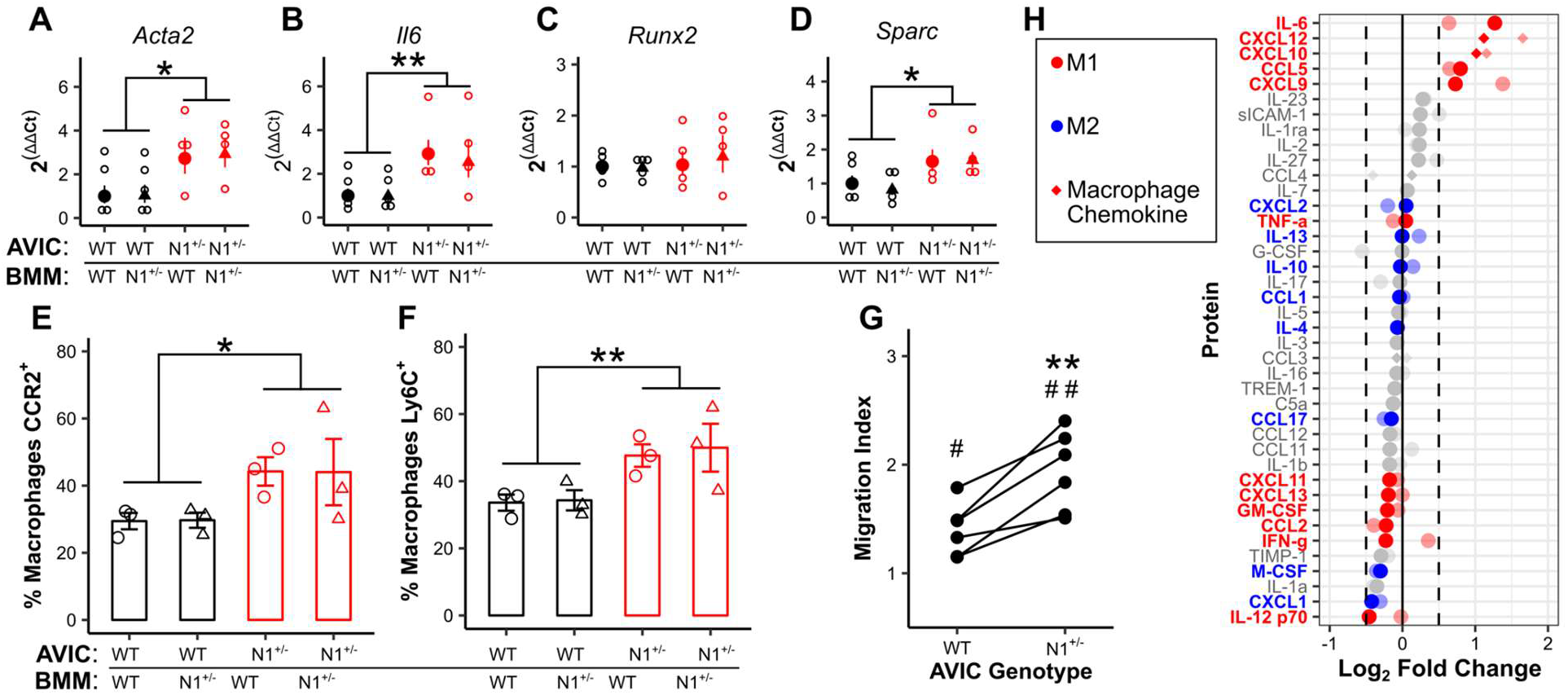
*Notch1*^+/-^ aortic valve interstitial cells promote calcification and macrophage maturation *in vitro* through altered cytokine secretion. Wild-type (WT) and *Notch1*^+/-^ (N1^+/-^) aortic valve interstitial cells (AVICs) were cultured with WT and N1^+/-^ bone marrow-derived macrophages (BMMs) and assayed for transcription of common markers of dystrophic (A, B) and osteogenic (C, D) calcification, regardless of BMM genotype. Flow cytometry was performed on BMMs for CCR2 (E) and Ly6C (F, black = WT AVICs; red = N1^+/-^ BMMs; circle = WT BMMs, triangle = N1^+/-^ BMMs). (G) Migration assays using N1^+/-^ and WT AVIC-cultured media and uncultured media controls. (H) Cytokine microarray analysis of secreted media from WT and N1^+/-^ AVICs. Data is plotted as fold change in N1^+/-^ AVIC-cultured media compared to WT control. Full color points are cultured media; faded points are cell lysates. Points represent an average of two experiments with 3 pooled samples in each. All summary data represent mean ± s.e.m. Data were analyzed by two-way ANOVA on untransformed ΔCt values (A-D) or flow populations (E, F), or one-way ANOVA followed by paired, two-tailed t tests with Holm-Sidak corrections (G). *P < 0.05 from wild-type AVICs, **P < 0.01 from wild-type AVICs, #P < 0.05 from monoculture control, ##P < 0.01 from monoculture control. N = biological replicates.

### Macrophages Promote Osteogenic, and not Dystrophic, Calcification of AVICs

Given the increased macrophage recruitment and polarization in the *Notch1*^+/-^ model, we sought to determine how macrophages affect AVIC calcification. When cultured with macrophages, AVICs formed more calcific nodules *in vitro* (Fig 4A). We then cultured AVICs either in monoculture (Mono), in Transwell culture with macrophages (TW), or in direct coculture with macrophages (CC) (Fig 4B). RNA and protein were then isolated from AVICs after MACS separation. RT-qPCR revealed increases in osteogenic calcification markers in both Transwell and, more profoundly, direct coculture (Fig 4C-E). There was no change in dystrophic calcification markers (Fig 4F-H). AVIC-specific expression was confirmed by immunofluorescent staining for CD68 (macrophages) and RUNX2 (Fig 4I). To further assess this relationship, macrophage proximity to RUNX2-positive and αSMA-positive AVICs was calculated (detailed methods in the supplemental material, Supp Fig II) (Fig 4J, K). RUNX2-positive AVICs (osteoblasts, osteogenic calcification) were closer to macrophages than expected, while αSMA-positive AVICs (myofibroblasts, dystrophic) were normally distributed as expected (Fig 4L). These data conclude that macrophages promote osteogenic, but not dystrophic, AVIC calcification.

**Figure 4.**
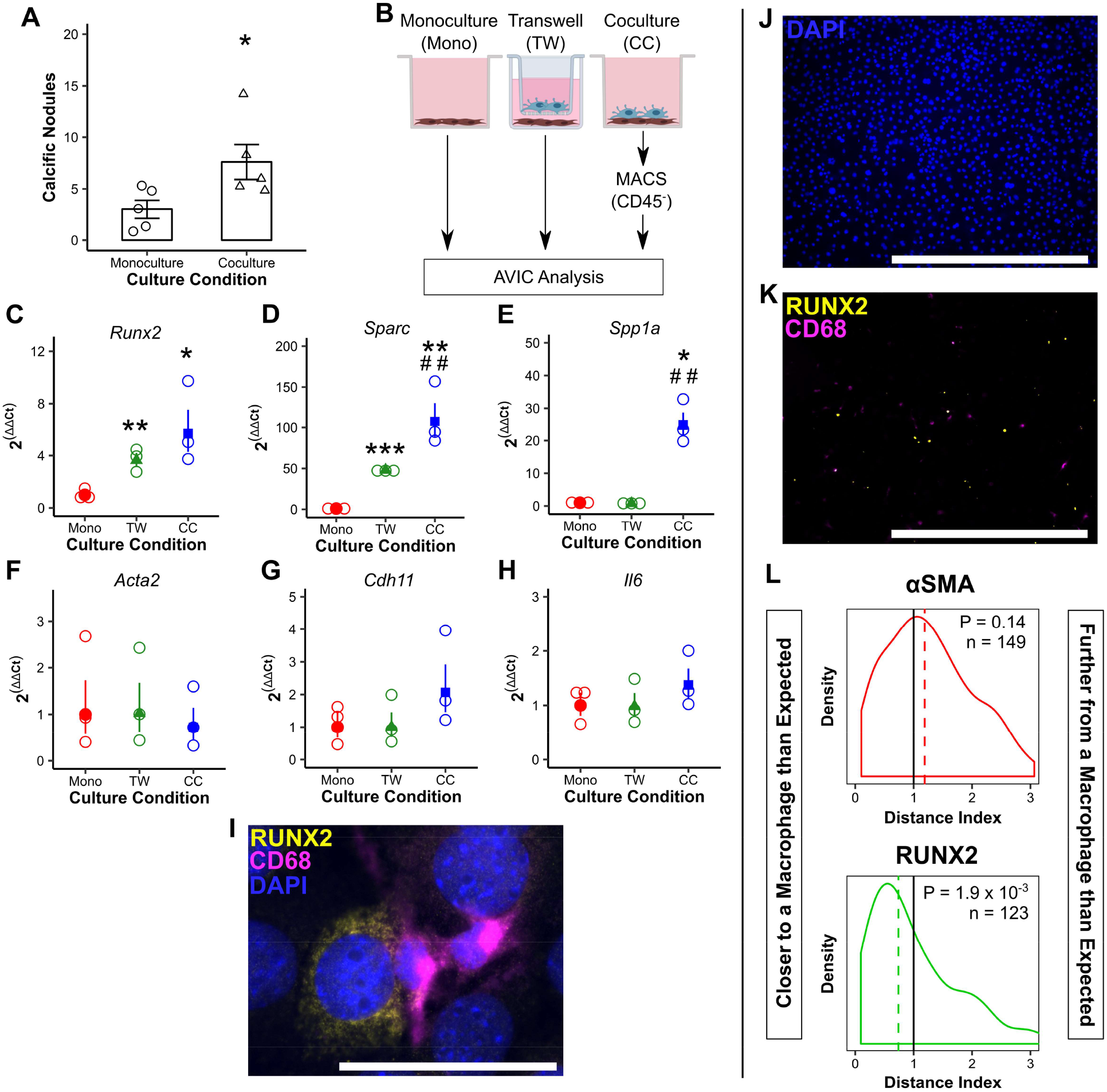
Macrophages promote osteogenic but not dystrophic calcification of aortic valve interstitial cells. Cocultures of aortic valve interstitial cells (AVICs) with bone marrow-derived macrophages (BMMs) were assayed for calcific nodule formation (A). AVICs cultured in monoculture (Mono), Transwell culture with BMMs (TW), or direct coculture with BMMs (CC) (B) were assayed for transcription of markers of osteogenic (C-E) and dystrophic (F-H) calcification. Images of cocultures stained for RUNX2, CD68, and DAPI (I-K) were analyzed by a Monte Carlo-assisted simulation to calculate expected distance and distance index between activated AVICs and BMMs (L). Scale bars = 50 µm (I) and 1 mm (J, K). All summary data represent mean ± s.e.m. Data were analyzed by Mann Whitney U test (A), one-way ANOVA followed by two-tailed t tests with Holm-Sidak corrections on untransformed ΔCt values (C-H), or one sample Wilcoxon Signed-Rank test on log-transformed data (L). *P < 0.05, **P < 0.01, ***P < 0.001 from monoculture AVICs; ##P < 0.01 from Transwell AVICs. N = biological replicates (B-K) or activated AVICs (L).

### Macrophages Promote Calcification through Altered STAT3 Splicing

In addition to canonical osteogenic signaling, we hypothesized that STAT3-mediated inflammation played a role in RUNX2 activation due to previous studies in CAVD,^24^ and the role of STAT3 in other fibrotic inflammatory diseases, including cardiovascular disease.^16–18^ AVICs cultured with macrophages had no increase in STAT3 phosphorylation or total STAT3 but did show a marked decrease in STAT3β expression (Fig 5A-E, Supp Fig VIII). STAT3β is an alternative splice product of the STAT3 gene that inhibits canonical STAT3 signaling mediated through STAT3α.^42, 43^ RT-qPCR of STAT3 transcriptional targets *Icam1* and *Vegfa* confirmed an increase in STAT3 activity (Fig 5F). To assess the role of STAT3β as an anti-calcification signaling molecule in human disease, excised AVs from patients undergoing AVR were analyzed. Tissue involved in disease had decreased STAT3β expression and increased RUNX2 expression compared to adjacent uninvolved tissue from the same patients (Fig 5G, H, Supp Fig IX, X). Across all samples, RUNX2 expression negatively correlated with the STAT3β fraction (Fig 5I).

**Figure 5.**
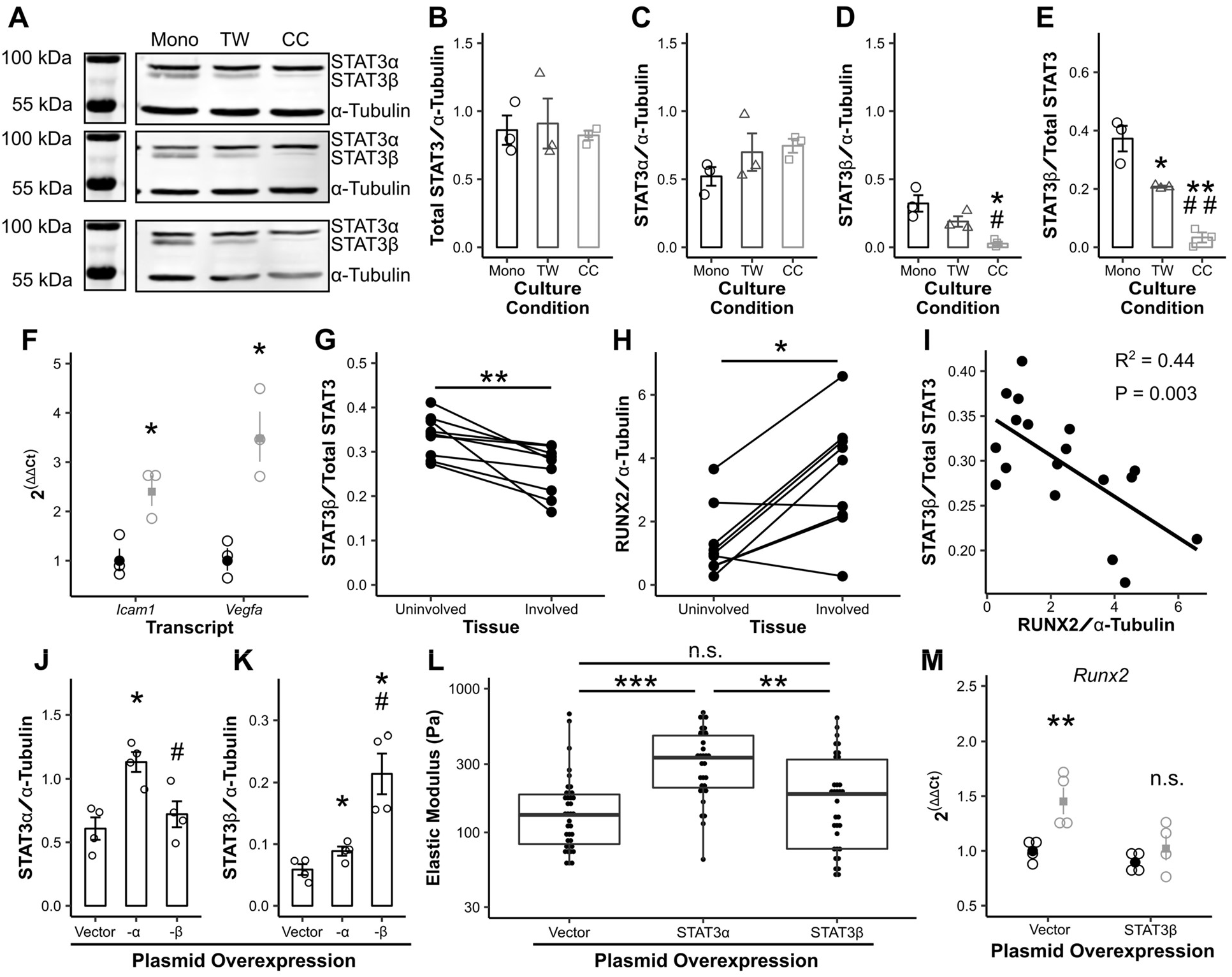
Macrophages promote osteogenic calcification of aortic valve interstitial cells through altered STAT3 splicing. Overall STAT3 expression (A, B), STAT3α expression (C), STAT3β expression (D, E), and expression of STAT3-associated transcripts (F) across 3 biological coculture replicates (F, black = monoculture [Mono], gray = Transwell [TW], light gray = coculture [CC]). STAT3β and RUNX2 expression was assayed by Western blot in calcified human aortic valves divided into involved and uninvolved tissue (G-I). Plasmid overexpression of STAT3α and β was performed (J, K), and cellular stiffness measured by micropipette (L). Overexpression of STAT3β was performed prior to coculture and cocultures assayed for Runx2 transcription. Bars and dot plots represent mean ± s.e.m. Boxplots represent 25^th^, 50^th^, and 75^th^ percentiles. Data were analyzed by one-way ANOVA followed by two-tailed t tests with Holm-Sidak corrections on densitometry data (B-E, J, K) or untransformed ΔCt values (F, M); paired Mann Whitney U tests (G, H); linear regression (I); or Kruskal-Wallis followed by Mann Whitney U tests with Holm-Sidak corrections (L). *P < 0.05, **P < 0.01, ***P < 0.001 from monoculture AVICs (D-F, M), uninvolved aortic valve tissue (G, H) or vector control (J, K); #P < 0.05, ##P < 0.01 from Transwell AVICs (D-E) or STAT3α transfection (J, K). N = biological replicates (B-K, M) or tests of individual cells across 3 biological replicates in 2 independent experiments (L).

Two STAT3 blockade strategies were assessed for efficacy in mitigating calcification. First, AVICs were treated with Stattic, a STAT3 phosphorylation inhibitor, and assayed for cellular stiffness and *Runx2* transcription. Stattic treatment decreased cellular stiffness but increased *Runx2* transcription in monoculture (Supp Fig XI). Separately, STAT3α and -β plasmids were used to artificially manipulate STAT3 splicing (Fig 5J, K, Supp Fig XII). Such transfections had no effect on calcification-associated transcripts in monoculture AVICs, but STAT3α overexpression increased cellular stiffness (Fig 5L). In the coculture model, STAT3β overexpression rescued *Runx2* transcription (Fig 5M).

## Discussion

While the cardiovascular immunology field has developed at a rapid pace, the role of immune cells in CAVD has remained unclear.^9, 44^ Here, we have focused on macrophages, which make up the majority of hematopoietic cells in the AV in both health and disease.^8^ We have shown that *Notch1*^+/-^ AVICs promote macrophage maturation and infiltration, and that macrophages promote AVIC calcification through STAT3-dependent mechanisms (Fig 6).

**Figure 6.**
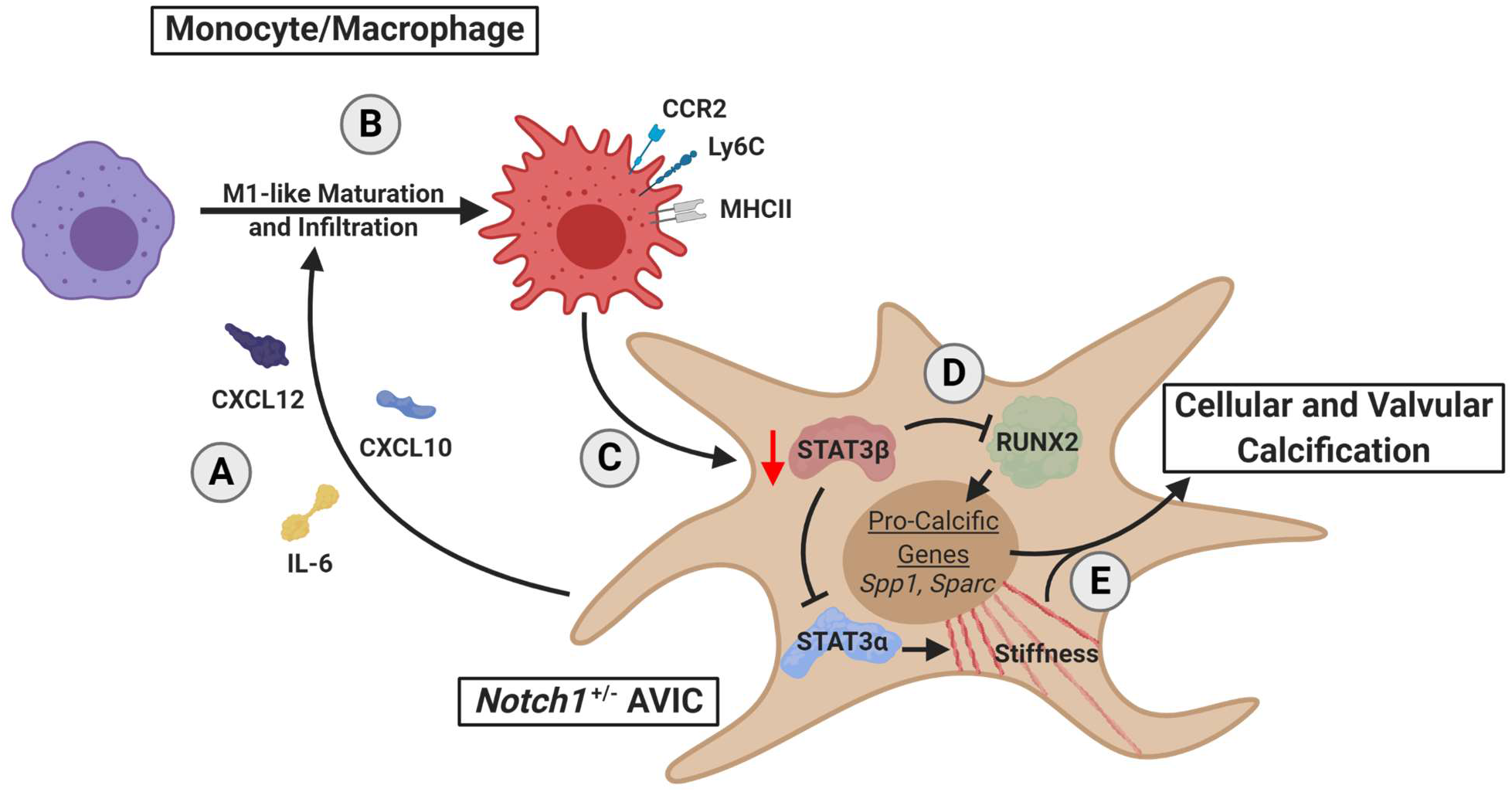
Proposed mechanism for macrophage-associated calcification in *Notch1*^+/-^ calcific aortic valve disease. *Notch1*^+/-^ aortic valve interstitial cells (AVICs) secrete pro-inflammatory factors (A) leading to increased macrophage infiltration and maturation to an M1-like phenotype (B). These infiltrating macrophages alter STAT3 splice products to decrease STAT3β (C). Decrease of STAT3β removes inhibition of STAT3α and RUNX2 (D), promoting cellular stiffening and expression of osteogenic transcripts, respectively, and leading to valvular calcification (E).

First, we used flow cytometry to show that the *Notch1*^+/-^ model has an increased myeloid compartment in the valve and increased MHCII positivity prior to disease progression (Fig 1). This aligns with previous data that murine aging and human disease correlate with M1 polarization,^5, 8, 45^ and that M1 polarization itself promotes cellular calcification.^5, 46^ This suggests that perhaps differences in macrophage phenotype at baseline in *Notch1*^+/-^ mice are driving AV phenotypes. In addition, *NOTCH1* is known to inhibit myeloid cell maturation, reflected in our data by an increased myeloid compartment in the *Notch1*^+/-^ valve.^30, 31^ Together, these data strengthened our hypothesis that altered hematopoietic cells drive disease in the *Notch1*^+/-^ model. Thus, we used bone marrow transplants to assess the role of the *Notch1*^+/-^ mutation in valve cells vs macrophages with increased MHCII positivity or other hematopoietic changes.

Bone marrow transplant experiments show that *Notch1*^+/-^ AV cells promote macrophage infiltration regardless of macrophage genotype (Fig 2). Contrary to our hypothesis, it seems that hematopoietic cells play their role in response to altered valve cell phenotypes. *Notch1*^+/-^ mice regardless of bone marrow genotype had increased macrophage infiltration, and infiltrating macrophages were more often MHCII^+^. Thus, *Notch1*^+/-^ valve cells are likely the driving force behind valve pathology by both driving traditional disease markers and recruiting hematopoietic cells that promote calcification. Interestingly, *Notch1*^+/-^ AVs had increased leaflet thickness and macrophage infiltration prior to development of hemodynamic obstruction (Supp Fig XIII). In combination with the data in Figure 1, this argues that macrophage infiltration is an early event in CAVD development, occurring before stenosis, rather than an anti-calcific compensation or a non-specific recruitment in response to fibrosis and ectopic calcification. The contribution of *Notch1* haploinsufficiency through AVICs rather than macrophages was replicated *in vitro* (Fig 3), reinforcing that *Notch1* AVICs rather than hematopoietic cells drive differences in calcification.

We built off of these findings with *in vitro* studies to explore how *Notch1*^+/-^ AVICs alter macrophage phenotypes and infiltration (Fig 3). *Notch1*^+/-^ AVICs promote increased migration and M1 polarization by flow cytometry when compared to media from wild-type AVICs. Reinforcing the importance of *Notch1* haploinsufficiency in AVICs specifically, *Notch1*^+/-^ macrophages responded similarly. Notably, while both experiments used the same antibody panel, the M1 phenotype induced *in vitro* is characterized by Ly6C and CCR2 positivity and no change in MHCII, while the *in vivo* data showed instead an increase in MHCII^+^ macrophages but no change in Ly6C or CCR2 by flow cytometry. It is possible that this difference is due to the timelines involved. High Ly6C expression defines inflammatory monocytes and macrophages,^47, 48^ and CCR2 is necessary for recruitment of such Ly6C^hi^ monocytes.^49, 50^ Thus, the roles of Ly6C and CCR2 are in the recruitment and egress of monocytes and macrophages, but expression is variable and can decrease after entering the tissue.^51, 52^ Alternatively, MHCII expression is a later development upon macrophage extravasation into the tissue that allows for antigen presentation and an adaptive immune response.^51, 53^ This would explain the increased MHCII expression seen in macrophages within murine valves.

Cytokine microarrays on the AVIC secreted media confirmed an increase in factors that induce migration and M1 polarization, which, in addition to our data showing increased prevalence in the *Notch1*^+/-^ model, has previously been shown to promote AVIC calcification.^5^ Together, these *in vivo* and *in vitro* phenomena provide a mechanism for macrophage involvement in *NOTCH1*-associated CAVD. They also highlight an additional lens for the interpretation of transcriptomic and proteomic datasets like that reported by Schlotter, et al.^54^ Such studies commonly consider how changes in the proteome of the valve or vascular compartment affect AVIC phenotypes. For example, Aguado, et al. have used the serum from patients undergoing transcatheter aortic valve replacement (TAVR) as a treatment and shown how differences in secreted factors affect AVIC calcification.^4^ Our results contribute to the body of literature suggesting that the effects of these secreted factors on immune cell recruitment and activation may also play a significant role in CAVD pathophysiology.

Our remaining studies turned to the question of how macrophages alter AVIC calcification. We utilized a Transwell model to show that not only does the macrophage secretome promote calcification, but that physical interaction increases this effect (Fig 4). This is perhaps due to a macrophage-to-AVIC signal, but considering the findings that AVICs promote macrophage activation, it is also possible that through physical interaction AVICs induced a further activated macrophage state wherein macrophages secreted additional pro-calcification cytokines. Nonetheless, macrophages clearly induced AVIC calcification in a proximity-dependent manner. Unintuitively, macrophages promoted osteogenic calcification and not dystrophic calcification, which is characterized by cytokine production and myofibroblast transition. To test the hypothesis that myofibroblast transition wasn’t increased *in toto*, but that myofibroblast activation was occurring closer to macrophages, we developed an image analysis algorithm. The results instead confirmed the above findings: αSMA-positive myofibroblasts were normally distributed around their expected distance, while RUNX2-positive osteoblast-like cells were significantly closer to macrophages than expected. This confirms the Transwell finding that proximity to macrophages is associated with increased osteogenic but not dystrophic calcification.

We then tested the hypothesis that STAT3 was mediating the connection between macrophage-secreted factors and RUNX2 expression. There was no increase in STAT3 phosphorylation or STAT3 expression on the whole, but a drastic shift in STAT3 splicing, decreasing the inhibitory STAT3β splice product and increasing the canonical STAT3α splice product (Fig 5). We confirmed an associated increase in canonical STAT3 signaling as measured through increased *Vegfa* and *Icam1* transcription: two signaling markers previously described in CAVD.^55, 56^ Importantly, STAT3β not only opposes STAT3α activity, thereby decreasing RUNX2 transcription, but also directly binds to RUNX2, preventing its activity as a transcription factor.^57^ These phenomena translated to human AVs: STAT3β decreases in calcified regions of diseased AVs and negatively correlates with RUNX2 expression. We then used overexpression models to manipulate STAT3 splicing directly. In AVIC monoculture, manipulation of STAT3 splicing ratios altered cellular stiffness, a calcification marker, and in coculture this manipulation mitigated increased RUNX2 transcription.

We attempted to understand the mechanism of STAT3β rescue by blocking STAT3 activity through dephosphorylation. This allowed us another angle through which to inhibit canonical STAT3 activity and test the hypothesis that STAT3β was acting primarily through STAT3α inhibition. Treatment with Stattic, a STAT3 phosphorylation inhibitor, decreased cellular stiffness but increased RUNX2 expression. This leads to the conclusion that STAT3β is functioning to inhibit calcification through its own unique characteristics, perhaps requiring phosphorylation, rather than solely through an auto-inhibitory function against canonical STAT3α signaling. The ability of STAT3β to bind RUNX2 and inhibit its function as a transcription factor may be a key step in its calcification-mitigating capabilities shown here.^57^

Increased study of immune cells in CAVD pathogenesis and pathophysiology provides an avenue for both pre- and post-AVR therapies. Unlike the native valve, bioprosthetic valves used in AVR do not replicate the typical AV microenvironment. Restenosis of these valves presents a challenge that likely could not be solved with a drug targeting native AVIC transformation. However, bioprosthetic restenosis does involve immune cell infiltrate.^12^ Thus, macrophage-associated calcification could be targeted both pre- and post-AVR. It is necessary to further probe the role of macrophages as CAVD initiators. Although others have depleted macrophages with liposomal clodronate, this experimental model has many off-target effects and is not completely reliable.^13^ Instead, a myeloid-targeting Cre or antibody depletion may be appropriate models for investigation.

It is crucial for those studying STAT3 biology to understand the inciting events underlying STAT3 splicing. Some explanations have been offered, one being decreased adenosine deaminase acting on RNA 1 (ADAR1) activity.^58^ We found opposite trends, with *Adar1* transcription increasing with decreased expression of STAT3β (Supp Fig XIV). Understanding the dynamic splicing of STAT3 would be impactful for many fields within biomedical science. Additional study of the STAT3 axis in CAVD murine models will shed further light on the utility of this targeting strategy.

### Limitations

We have used primarily murine data throughout this manuscript. This has allowed us both to study the *Notch1*^+/-^ CAVD model, and to use coculture models with syngeneic macrophages. However, we have not replicated all findings with human samples. Second, it is possible in this murine model that there are resident hematopoietic progenitor cells in the AV that have persisted through irradiation and proliferated. However, literature in this mouse model have shown that all hematopoietic cells in the valve are perpetually recruited, rather than existing as resident cells,^59^ and we have used a high radiation dose to minimize this risk. Finally, the murine model of CAVD used here is subject to relatively large variance, making some studies underpowered for phenotype detection by echocardiography. Thus, we have focused on quantitative histological and immunofluorescence methods that we believe capture with integrity the extent of disease in mice.

## Conclusions

Herein, we report heightened macrophage infiltration and polarization in *NOTCH1*-associated CAVD, driven by altered cytokine secretion of AVICs. This increased interaction between macrophages and AVICs promotes AVIC calcification in part through STAT3 splicing. Altered STAT3 splicing is found in calcified human AVs, and splicing manipulation opposes macrophage-induced calcification. These findings suggest that cellular inflammation and the STAT3 axis may play a targetable role in CAVD.

## Acknowledgments

The authors would like to thank Caleb Snider for technical assistance with murine irradiation and Ethan Joll for discussion of image processing and data analysis.

## Nonstandard Abbreviations and Acronyms

αSMA: alpha smooth muscle actin
AV: aortic valve
AVIC: aortic valve interstitial cell
AVR: aortic valve replacement
BMM: bone marrow-derived macrophage
CAVD: calcific aortic valve disease
IL: interleukin
MACS: magnetic-assisted cell sorting
MHCII: major histocompatibility complex II
N1^+/-^: *Notch1*^+/-^
RT-qPCR: quantitative real time-polymerase chain reaction
STAT3: signal trasducer and activator of transcription 3
TAVR: transcatheter aortic valve replacement
TGF-β1: transforming growth factor beta 1
TNFα: tumor necrosis factor alpha
WT: wild-type

## Sources of Funding

This work was funded by the National Institutes of Health (F30-HL147464, R35-HL135790, and T32-GM007347) and Fondation Leducq.

## Disclosures

Dr. Lindman has received research grants from Edwards Lifesciences and Roche Diagnostics; served on scientific advisory boards for Roche Diagnostics; and has been a consultant to Medtronic and Roche Diagnostics.

## Highlights

- *Notch1*^+/-^ valve disease involves increased inflammatory macrophage infiltration, driven by altered cytokine secretion of aortic valve interstitial cells.
- Macrophages promote osteogenic, but not dystrophic, calcification of aortic valve interstitial cells.
- Macrophages promote calcification in part through altered STAT3 splicing leading to a decrease in the STAT3β splice product.

