## Supplemental for "Macrophages Promote Aortic Valve Cell Calcification Through STAT3 Splicing"

### Supplemental Methods

Image Analysis

In order to determine if activated AVICs were distributed unevenly throughout the coculture landscape within any given x-y field of view, an image processing algorithm was developed to test the hypothesis that activated cells were more likely to be near macrophages. After staining, images in each channel were blurred using a Gaussian filter with a standard deviation of 4 and thresholded by Otsu's thresholding method. This mask generated positive regions, which were gated by size to identify individual cells. This process was performed in each channel to identify $CD68^+$ macrophages, and either $RUNX2^+$ or $\alpha SMA^+$ activated AVICs.

In every image, the centroid of each activated AVIC was determined and a distance index was defined. In particular, for each identified AVIC, the location was compared to the location of each of the *n* identified macrophages and the distance to the nearest macrophage was recorded. All distances ≤ 10 μm were removed from the analysis to account for mistaken identification of one cell as both an AVIC and macrophage. Sensitivity analyses confirmed that this did not affect the conclusion. In order to determine the "expected distance" from the activated AVIC to the nearest macrophage, *N* random macrophages were placed across the x and y axes of the image and the distance from the current AVIC to the nearest random macrophage location was recorded. This randomized process was repeated 500 times in a Monte Carlo simulation, and the median distance to the nearest macrophage was recorded as the "expected distance" to the nearest macrophage. At that point, the real distance to the nearest macrophage was divided by the expected distance and recorded as the distance index of the activated AVIC. A density plot of the distance index of all activated AVICs is shown in Figure 5. The R package '*EBImage*' was used for all image processing.

Micropipette Aspiration

Capillary tubes (World Precision Instruments, Sarasota, FL) were coated with Sigmacote (MilliporeSigma, St. Louis, MO), sterilized with 70% ethanol, and allowed to dry. Coated tubes were then pulled with a P-97 micropipette puller (Sutter Instrument, Novato, CA), fractured with an MF-1 microforge (Technical Products International, St. Louis, MO) to an internal diameter of approximately 6 μm, and bent to an angle allowing for the micropipette to lie parallel to the plate upon use. Pressures were applied using a custom-built pressure regulator system with an MCFS-EZ microfluidics controller (Fluigent, Le Kremlin-Bicêtre, France).

Following treatment, AVICs were lifted with Accutase, resuspended in 500 μL MACS buffer, and kept on ice until use. Aspiration was performed on at least 10 cells from each condition and biological replicate each day. Tests were performed by linearly increasing the applied suction pressure by 8 Pa/s over 60 seconds to a final aspiration pressure of 0.48 kPa. The aspirated length of each cell was measured manually from video recorded at a rate of 2 frames/s using a microscope-mounted camera.

After all data was recorded, aspirated length of each cell was measured manually and the effective stiffness (*E*) was determined using a half-space elastic model given below:

$$E = \varphi n \left(\frac{3r}{2\pi}\right)\left(\frac{\Delta P}{L}\right)$$

where $\varphi(\eta)$ is the wall function and is equal to 2.1 (dimensionless parameter calculated from the ratio of the pipette inner radius to the wall thickness), *r* is the micropipette inner radius, and *ΔP/L* is the slope of the linear applied pressure vs. aspirated cell length curve.

**Supplemental Figures**

Supplemental Figure I. Confirmation of bone marrow-derived macrophage phenotype in coculture experiments.

Supplemental Figure II. Image proximity analysis workflow.

Supplemental Figure III. Raw micropipette analysis data.

Supplemental Figure IV. Wild-type and *Notch1*$^{+/-}$ bone marrow-derived macrophages.

Supplemental Figure V. MHCII$^{-}$ macrophages in wild-type and *Notch1*$^{+/-}$ valves.

Supplemental Figure VI. *Notch1*$^{+/-}$ macrophage migration towards AVIC-secreted media.

Supplemental Figure VII. Raw Proteome Profiler microarray results.

Supplemental Figure VIII. STAT3 splicing in AVICs exposed to macrophages.

Supplemental Figure IX. STAT3 splicing in human calcific aortic valve disease.

Supplemental Figure X. RUNX2 in human calcific aortic valve disease.

Supplemental Figure XI. Stattic treatment of AVICs.

Supplemental Figure XII. STAT3 plasmid transfection.

Supplemental Figure XIII. Echocardiography metrics in wild-type and *Notch1*$^{+/-}$ mice.

Supplemental Figure XIV. *Adar1* transcription in cocultured AVICs.

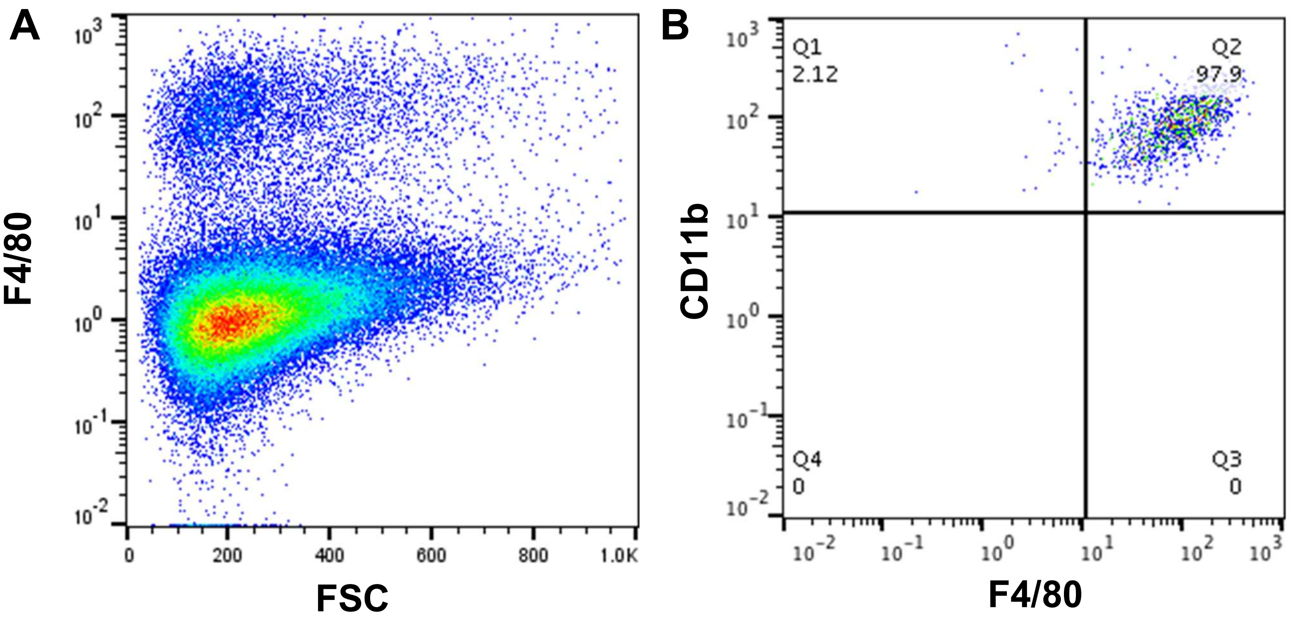


**Supplemental Figure I. Confirmation of bone marrow-derived macrophage phenotype in coculture experiments.** The F4/80$^{hi}$ macrophage population is seen in AVIC-macrophage coculture (A, representative plot). Among CD45$^{+}$ cells, >96% are CD11b$^{+}$ and F4/80$^{hi}$ in each of four biological replicates (B, representative plot).

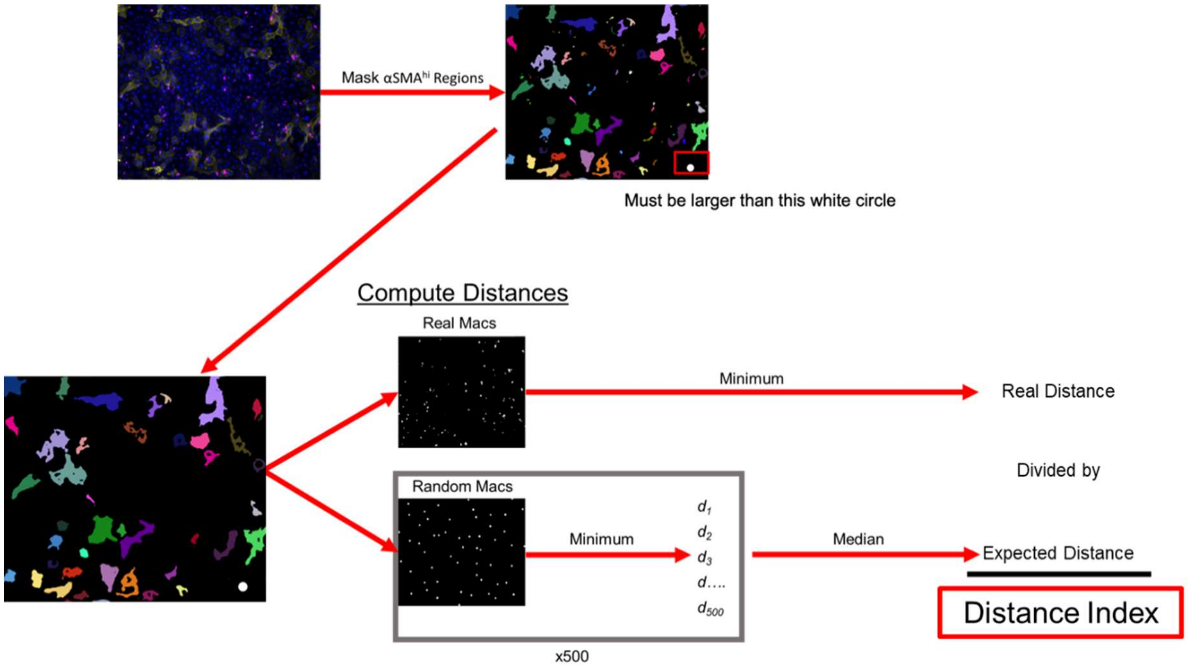


**Supplemental Figure II. Image proximity analysis workflow.** Images were masked for activated AVICs by RUNX2 or αSMA staining and real and expected distance to the nearest macrophage calculated. Additional details are included above in the supplemental methods.

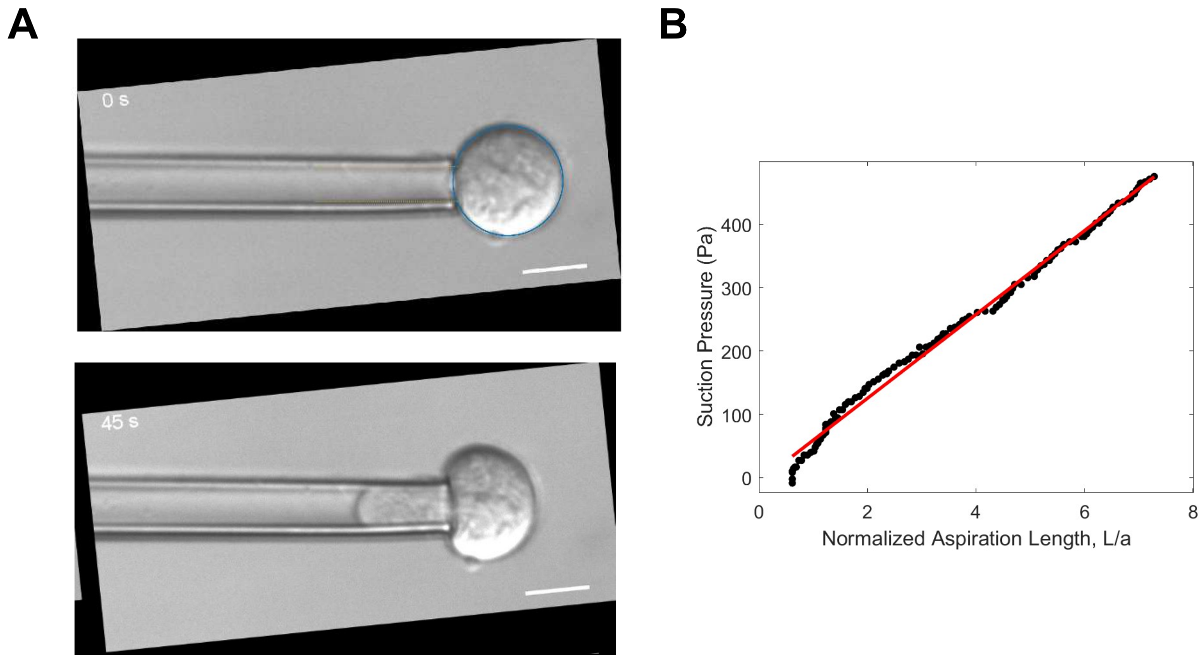


**Supplemental Figure III. Raw micropipette analysis data.** Stabilized images of cell aspiration were recorded (A) followed by measurement of the slope of suction pressure over normalized aspiration length to determine cellular elastic modulus (B).

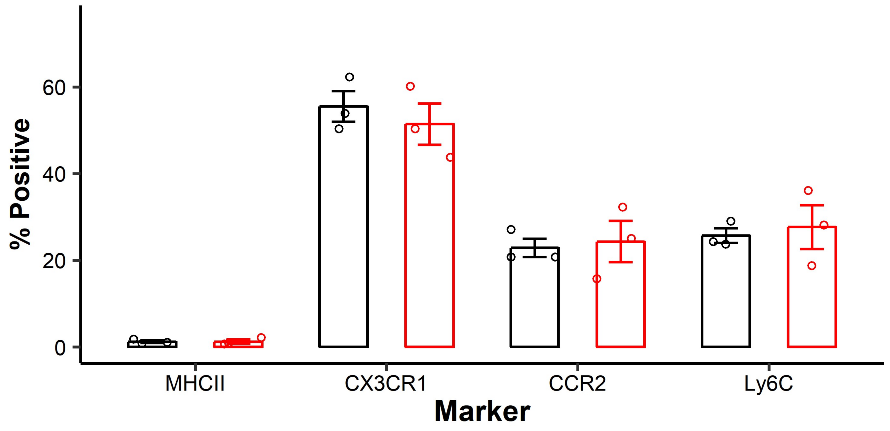


**Supplemental Figure IV. Wild-type and *Notch1*$^{+/-}$ bone marrow-derived macrophages.** Bone marrow-derived macrophages from wild-type (black) and *Notch1*$^{+/-}$ (red) mice have no differences in various markers of maturity. Bars represent mean ± s.e.m. All data analyzed by two-tailed *t* test.

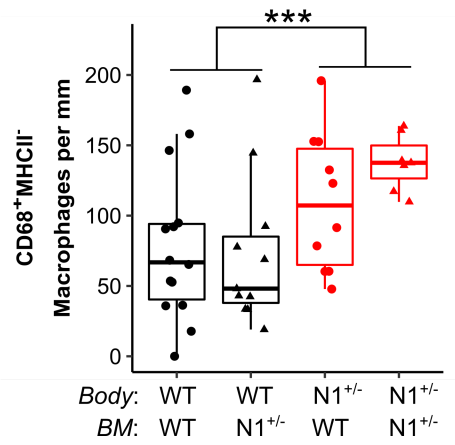


**Supplemental Figure V. MHCII$^{-}$ macrophages in wild-type and *Notch1*$^{+/-}$ valves.** There is an increase in MHCII$^{-}$ macrophages in *Notch1*$^{+/-}$ mice regardless of bone marrow genotype. Boxplots display the 25$^{th}$, 50$^{th}$, and 75$^{th}$ percentiles. Data were analyzed by two-way aligned rank transformed ANOVA. ***P < 0.001. N = biological replicates.

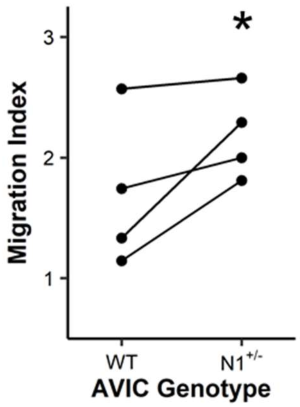


**Supplemental Figure VI. *Notch1*$^{+/-}$ macrophage migration towards AVIC-secreted media.** *Notch1*$^{+/-}$ AVIC-cultured media promotes macrophage migration compared to wild-type AVIC-cultured media or uncultured media control. Data were analyzed by one-way ANOVA followed by paired, two-tailed *t* tests with Holm-Sidak corrections (G). *P < 0.05 from wild-type AVICs, N = biological replicates.

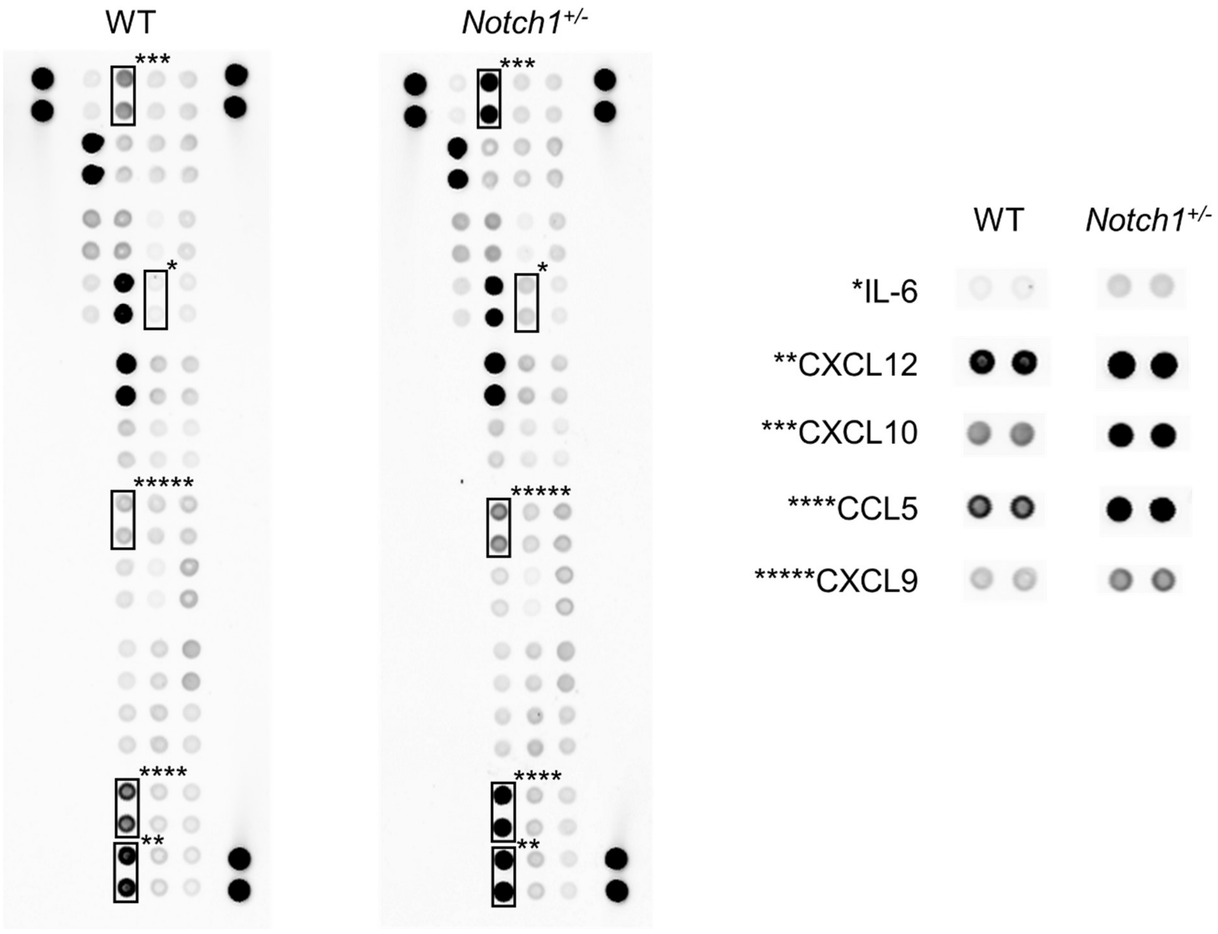


**Supplemental Figure VII. Raw Proteome Profiler microarray results.** Microarray of secreted factors from wild-type (WT) and *Notch1*$^{+/-}$ AVICs. *Denotes corresponding microarray spots cropped for comparison.

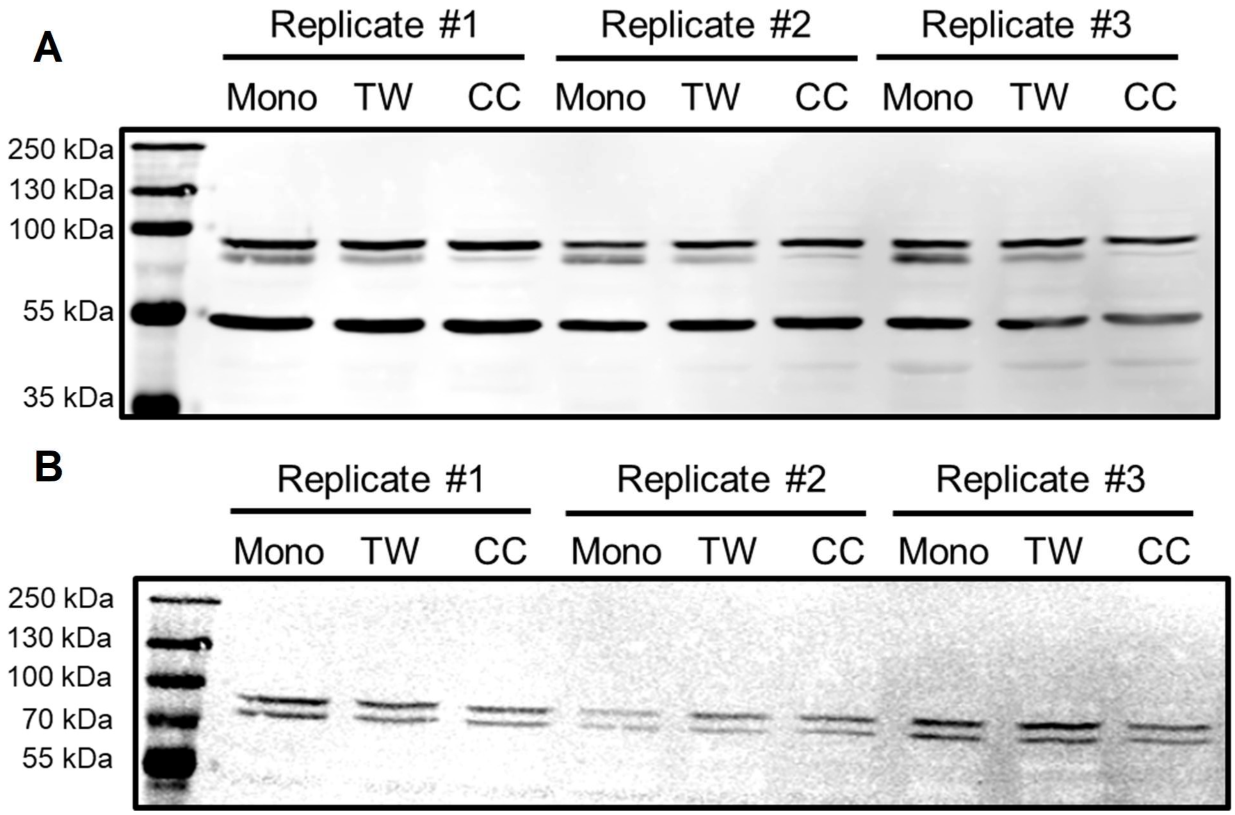


**Supplemental Figure VIII. STAT3 splicing in AVICs exposed to macrophages.** Raw Western blot data for STAT3 splicing (A) and phosphorylation (B) in AVICs in monoculture (Mono), Transwell culture (TW), or direct coculture (CC) with macrophages. p/STAT3α is stained at ~88 kDa with p/STAT3β just below. Loading control is α-Tubulin stained at ~50 kDa (A). STAT3 is visualized with anti-mouse IgG2a secondary antibody in the 700 channel, pSTAT3 with anti-rabbit IgG secondary antibody in the 800 channel, and α-Tubulin with anti-mouse IgG1 secondary antibody in the 700 channel.

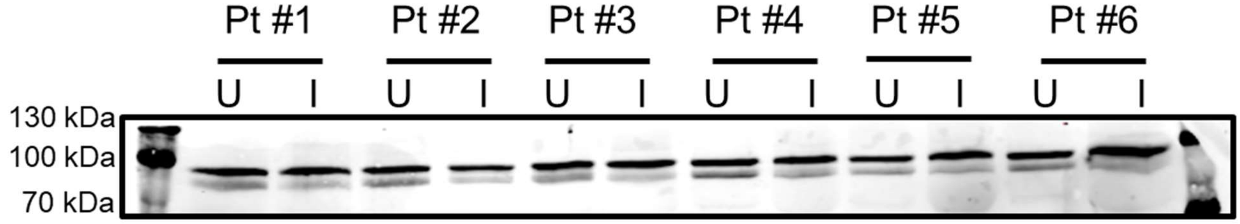


**Supplemental Figure IX. STAT3 splicing in human calcific aortic valve disease.** Representative raw Western blot data for STAT3 splicing in uninvolved (U) and involved (I) tissue from patients with calcific aortic valve disease. STAT3α is stained at ~88 kDa with STAT3β just below. Total STAT3 quantified from previous Western blot was used as loading control in order to normalize STAT3β to total STAT3. STAT3 is visualized with anti-mouse IgG2a secondary antibody in the 700 channel.

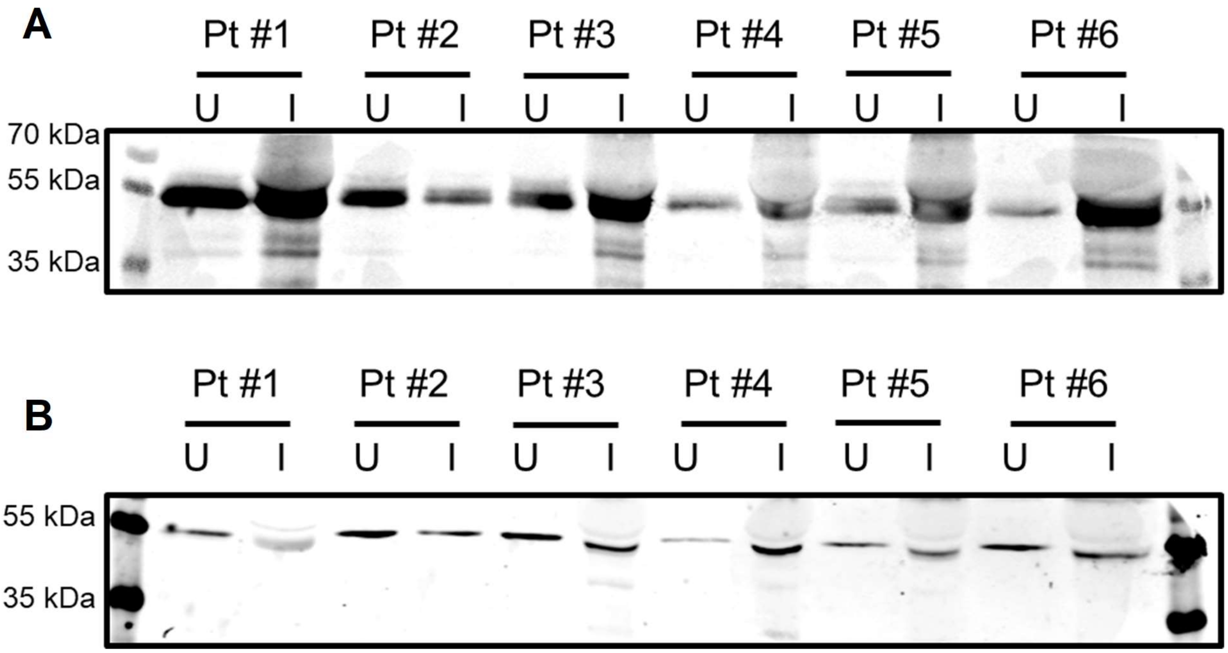


**Supplemental Figure X. RUNX2 in human calcific aortic valve disease.** Representative raw Western blot data for RUNX2 expression in uninvolved (U) and involved (I) tissue from patients with calcific aortic valve disease. RUNX2 is stained at ~56 kDa (A). Loading control is α-Tubulin stained at ~50 kDa (B). RUNX2 is visualized with anti-rabbit IgG1 secondary antibody in the 800 channel and α-Tubulin is visualized with anti-mouse IgG1 secondary antibody in the 700 channel. Although they are in separate channels, RUNX2 was stained first followed by α-Tubulin to prevent any bleedover of α-Tubulin signal into RUNX2 densitometry quantification.

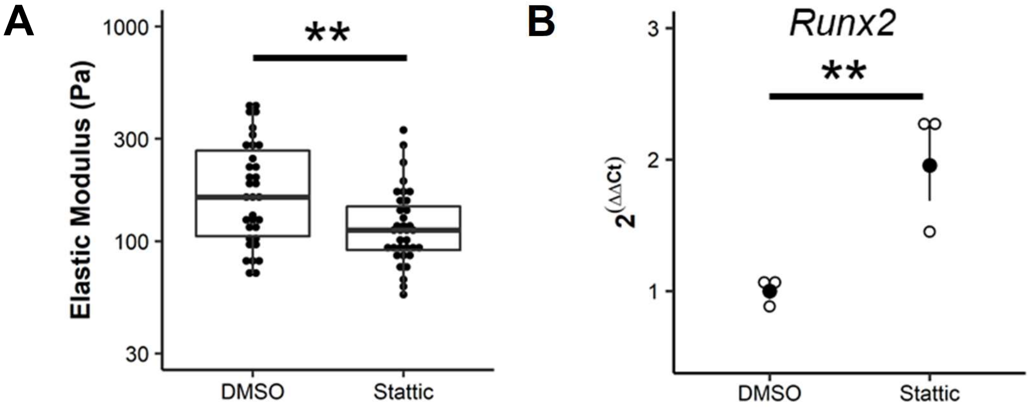


**Supplemental Figure XI. Stattic treatment of AVICs.** Treatment with 10 μM Stattic for two hours decreases cellular stiffness (A) but increases *Runx2* transcription measured after 10 additional hours in complete DMEM media (B). Boxplots display the 25$^{th}$, 50$^{th}$, and 75$^{th}$ percentiles. Summary data represent the mean ± s.e.m. (B). **P < 0.01 by Mann Whitney *U* test (A) or two-tailed *t* test on untransformed ΔCt values (B).

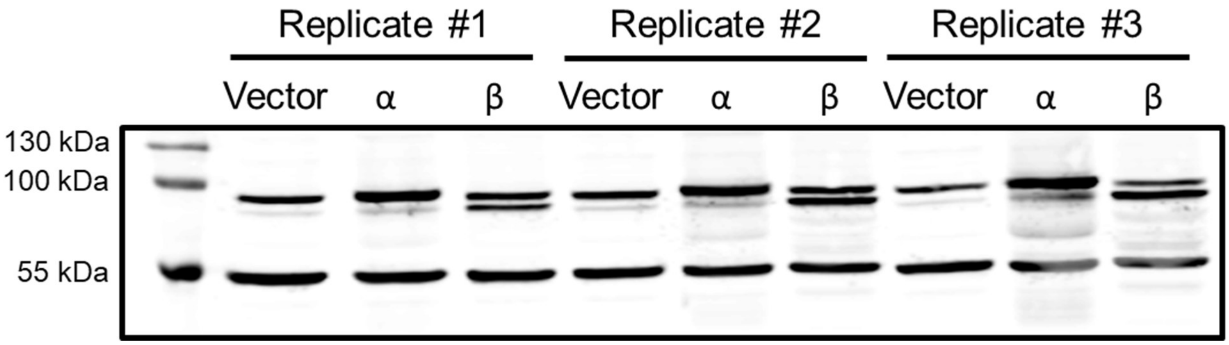


**Supplemental Figure XII. STAT3 plasmid transfection.** Representative raw Western blot data for STAT3 expression in samples transfected with empty vector plasmid (Vector), STAT3α overexpression plasmid (α), or STAT3β overexpression plasmid (β). STAT3α is stained at ~88 kDa with STAT3β just below. Loading control is α-Tubulin stained at ~50 kDa. STAT3 is visualized with anti-mouse IgG2a secondary antibody and α-Tubulin is visualized with anti-mouse IgG1 secondary antibody: both in the 700 channel.

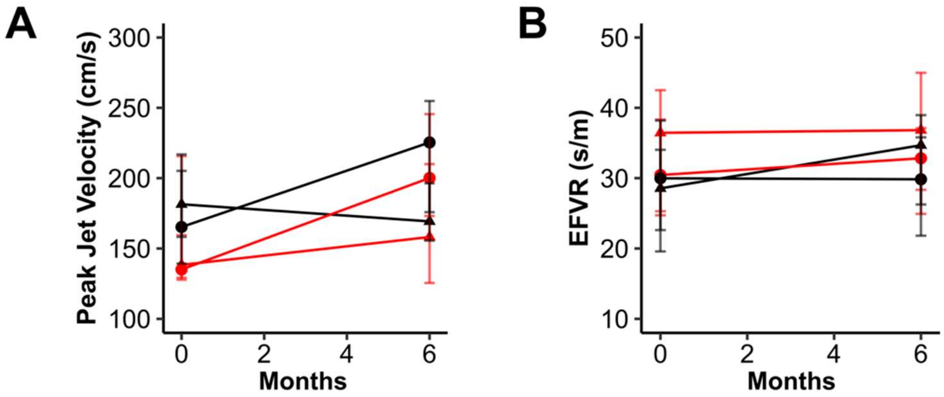


**Supplemental Figure XIII. Echocardiography metrics in wild-type and *Notch1*$^{+/-}$ mice.** Wild-type (black) and *Notch1*$^{+/-}$ (red) mice with wild-type (circle) and *Notch1*$^{+/-}$ (triangle) bone marrow have no differences in peak jet velocity (A) or ejection fraction-velocity ratio (EFVR) (B).

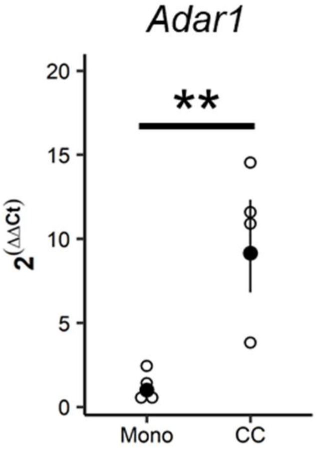


**Supplemental Figure XIV. *Adar1* transcription in cocultured AVICs.** Coculture of AVICs with macrophages increases transcription of *Adar1*. Summary data represent the mean ± s.e.m. **P < 0.01 by two-tailed *t* test.

### Major Resources Tables

#### Animals (in vivo studies)

| Species | Vendor or Source | Background Strain | Sex |
|---|---|---|---|
| *Mus musculus* | The Jackson Laboratory | C57BL/6J; *Notch1*$^{+/-}$ | 69 x M (BMT, BMM, FC, LSM)<br>45 x F (BMT, BMM) |

#### Animal breeding

| | Species | Vendor or Source | Background Strain | Other Information |
|---|---|---|---|---|
| **Parent** | *Mus musculus* | The Jackson Laboratory, Bar Harbor, ME | C57BL/6J | Breedings were between *Notch1*$^{+/-}$ and *Notch1*$^{+/+}$ animals backcrossed onto a C57BL/6J background to produce wild-type and *Notch1*$^{+/-}$ littermate controls |
| **Parent** | *Mus musculus* | Merryman Lab from Rossant Lab, University of Toronto | C57BL/6J; *Notch1*$^{+/-}$ | Breedings were between *Notch1*$^{+/-}$ and *Notch1*$^{+/+}$ animals on a C57BL/6J background to produce wild-type and *Notch1*$^{+/-}$ littermate controls |

#### Antibodies

| Target antigen | Vendor or Source | Catalog # | Working concentration | Lot # (preferred but not required) |
|---|---|---|---|---|
| αSMA-Cy5.5 | MilliporeSigma | C6198 | IF(1:300) LSM(3μg) | 058M4761V |
| α-Tubulin | Vanderbilt Molecular Biology Core | n/a | WB(1:1000) | n/a |
| CCR2-PE | BioLegend | 150609 | FC(1:50) | B278733 |
| CD11b-e450 | Thermo Fisher | 48-0112-82 | FC(1:400) | 4329941 |
| CD45-BV510 | BD Biosciences | 563891 | FC(1:800) | 9066967 |
| CD68-AF647 | Santa Cruz Biotechnology | sc-20060 | LSM(3μg) | |
| CD68-AF594 | BioLegend | 137020 | IF(1:200) | B239125 |
| CX3CR1-PerCP/Cy5.5 | BioLegend | 149009 | FC(1:250) | B271940 |
| F4/80-PE/Cy7 | BioLegend | 123114 | FC(1:400) | B265636 |
| Ly6C-FITC | BioLegend | 128006 | FC(1:700) | B270133 |
| MHCII-APC | BioLegend | 107614 | FC(1:1600) | B255462 |
| MHCII-FITC | Thermo Fisher | 11-5321-82 | IF(1:100) | 4322171 |
| RUNX2 | Novus Biologicals | NBP1-77461 | IF(1:100) | B-1 |
| Rabbit IgG-FITC | Abcam | ab6717 | IF(1:300) | 731506 |
| RUNX2 | Cell Signaling | 12556S | WB(1:1000) | 2 |
| STAT3 | Cell Signaling | 9139S | WB(1:1000) | 12 |
| pSTAT3 (Y705) | Cell Signaling | 9145S | WB(1:2000) | 34 |

#### Cultured Cells

| Name | Vendor or Source | Sex (F, M, or unknown) |
|---|---|---|
| Immortalized wild-type AVICs | Immorto Mice | 2 x M, 3 x F |
| Immortalized *Notch1*$^{+/-}$ AVICs | Immorto Mice | 2 x M, 2 x F |
| Bone marrow-derived macrophages | Wild-type and *Notch1*$^{+/-}$ mice | 12 x M, 12 x F |
